# Phenotypic Shifts in Neurons of the Dorsal Motor Nucleus of the Vagus Resulting from Chronic Cardiac Ischemia

**DOI:** 10.1101/2020.03.11.987982

**Authors:** Jonathan Gorky, Rajanikanth Vadigepalli, James Schwaber

**Affiliations:** Daniel Baugh Institute for Functional Genomics and Computational Biology Thomas Jefferson University, Philadelphia, PA, USA

## Abstract

Heart disease remains the number one cause of mortality in the world in spite of significant efforts aimed at treatment. The use of vagal stimulation in the treatment of heart failure has shown mixed successes (Dicarlo et al. 2013; Zannad et al. 2015), suggesting that the treatment has potential, but that the mechanism incompletely understood. Vagal activity confers a robust cardioprotective effect in both humans and animal models, preferentially originating specifically from the dorsal motor nucleus of the vagus (DMV) and deriving a significant benefit from intact gut projections, not just cardiac (Shinlapawittayatorn et al. 2013; Mastitskaya et al. 2012; Basalay et al. 2012). In order to examine the DMV response to heart failure, myocardial infarction was induced in male Sprague Dawley rats. DMV neurons were isolated in small pools of single cells using laser capture microdissection 1 week and 3 weeks after infarction and their gene expression assayed. The results show a transcriptional shift towards a neurosecretory phenotype starting at 1 week and increasing in recruitment of neurons to 3 weeks. The LAD ligation shift appears mediated in part by upregulation of Pax4a, a transcription factor most active during stem cell development of neurosecretory cells during embryonic development. This phenotype is characterized by upregulation of Cacna1d (Cav1.3) and Hcn2 along with increased expression of Cck and Sst. This work suggests that the neurons of the DMV adaptively respond to the dynamics present in the periphery, elucidating the means by which the nature of vagal activity responds to heart failure.

**Significance Statement:** The autonomic nervous system plays a significant role in the pathogenesis of cardiovascular disease. Through demonstration of shifting neuronal phenotypes in central neurons in response to peripheral stimuli, we suggest that neuron peptide or neurotransmitter phenotypes are not static in adult rodents. This suggests that even “reflexes” are modifiable dynamic systems. With such plasticity in the transcriptional programming existing in autonomic brain regions there can be new potential therapeutic interventions for cardiovascular disease aimed at leveraging the autonomic nervous system.

## Introduction

Heart failure continues to challenge the medical system with over 50% of patients dying within 5 years of diagnosis and 90% within 10 years (Go et al. 2013; Roger 2013). The association between heart failure and autonomic imbalance has been well-documented over the last several decades and it was hoped that restoration of balance through stimulation of the vagus nerve would prove therapeutic (Thayer et al. 2010; Sun et al. 2015; Pal et al. 2013; Floras and Ponikowski 2015; Jankowska et al. 2006; Ponikowski et al. 1997; Binkley et al. 1991). However, such treatments have proven to have inconsistent efficacy, sparking a new interest in understanding what exactly was being stimulated in efficacious instances and what about the context of the system permits a beneficial effect. It is not just a question of the effects of stimulating the afferent versus efferent vagal fibers, but also specifically which efferent projections among a large range of vagal cardiac effects. The vagal efferents are comprised of preganglionic neurons from two distinct regions of the medulla, the nucleus ambiguus (NA) and the dorsal motor nucleus of the vagus (DMV). The DMV projections differ from the NA projections in several ways, including which organs they project to as well as which postganglionic neurons in particular they synapse with at the heart (Jankowska et al. 2006).

The dorsal motor nucleus of the vagus contains vagal motor neurons that predominantly project to the gut (stomach, small intestine, proximal colon), but also project to the liver, pancreas, spleen, lungs, and heart (Jarvinen and Powley 1999). The DMV projections to the heart appear to have a relatively small role in the rhythmic regulation in the baro- and chemo-reflexes and in sinus arrhythmia coordinating with respiration (McAllen and Spyer 1978; McAllen et al. 2011; Jones et al. 1995; Jones et al. 1998). While it is clear that both the DMV and the NA can have profound cardiac effects and must in some way be coordinated in vagal influence on the heart, the lack of understanding of normal physiological effect have led to a diminished interest in research until recently when the phenomenon of cardioprotection from ischemic preconditioning was shown to rely upon vagal projections from the DMV and not the NA. This effect, in rats, has subsequently been determined to have a significant component that derives from the posterior gastric branch of the vagus, which originates entirely within the right side of the DMV (Mastitskaya et al. 2016; Fox and Powley 1985). This may have much broader implications into how the DMV responds to specific stressors to protect vital organs against potential subsequent stressors. Thus, in addition to direct vagal influences on the heart the vagus may also influence the heart indirectly. Recently, the role of the DMV in conferring a cardioprotective effect during the acute phase of cardiac ischemia-reperfusion injury has been described and validated in small animal models (Basalay et al. 2012). This effect appears to be at least partially mediated by DMV projections to the gut through release of a cardioprotective neuropeptides from the gut including glucagon-like peptide 1 and cholecystokinin among others (Mastitskaya et al. 2016; Basalay et al. 2016). These findings coupled with well-established links between overall gut health and cardiovascular disease support a significant role for the DMV in the response to heart failure (Cavallari et al. 2018; Kachur et al. 2018). In this context there is a foundation for the case that the gut has substantial influence on heart health lending it robustness to injury, with the DMV being the primary effector for top-down gut modulation.

In this work, we examine the shifts in gene expression and subsequent gene regulatory networks in a large number of neurons isolated from the DMV in animals subjected to left anterior descending artery (LAD) ligation and subsequent chronic ischemia. The results suggest a dramatic shift to a neurosecretory, enteroendocrine-like phenotype in a highly coordinated fashion in unique response to 3 weeks of chronic cardiac ischemia.

## Methods

### In vivo manipulations

All experiments were performed with the approval of the Institutional Animal Care and Use Committee at Thomas Jefferson University. All procedures and sacrifices were done within the same four hour circadian period. Male Sprague-Dawley rats (250-300g) were anesthetized with ketamine (100mg/kg) and xylazine (10mg/kg) and intubated with an 18g venous catheter and the animal was ventilated with oxygen-enriched room air using a rodent ventilator (Harvard Apparatus). A left thoracotomy was performed using aseptic technique and the heart was exteriorized. The left anterior descending (LAD) artery ligation was performed by passing a 6-0 prolene cardiac suture around the LAD artery and tying it off. Paling of the myocardium was observed before the heart was replaced back into the thoracic cavity and the thoracotomy closed with 4-0 silk suture. The sham ligation procedure involved the same preparations and left thoracotomy, but instead involved passing the 6-0 suture and needle around the LAD artery without tying the suture. The animals were then extubated and allowed to recover on a heating pad for four hours before being transported back to the holding facility. Animals were monitored post-surgically every 24 hours for signs of distress or improper healing and removed from the study if any such signs presented. After sacrifice, the hearts were collected and examined histologically to validate that the left ventricular remodeling did occur as expected (Figure 1).

**Figure 1:**
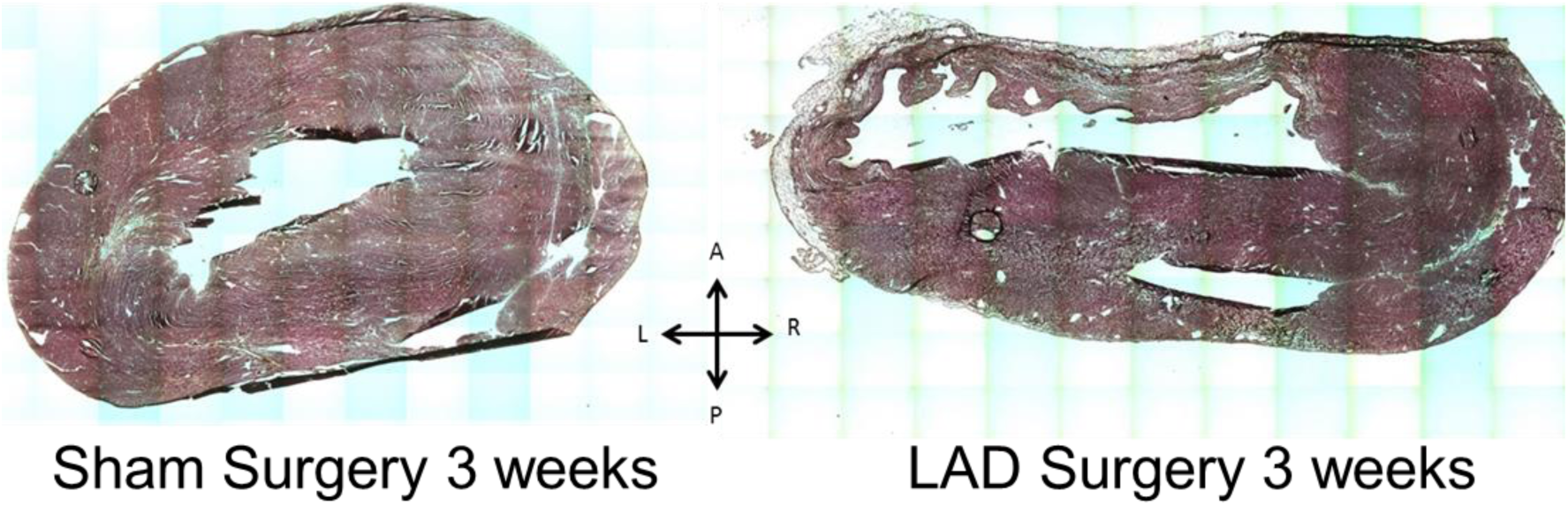
H & E staining of horizontal sections of example hearts validating the LAD ligation and sham ligation procedures anatomically. The left anterior wall of the left ventricle in the LAD ligation heart shows clear signs of myocardial atrophy and scarring that is absent in the sham ligation heart.

### Sample acquisition

After the appropriate survival period of either one week or three weeks, each animal sacrificed by rapid decapitation that was preceded by brief 60 seconds of 5% isoflurane in O_2_. The brain was quickly removed and placed into ice cold artificial cerebrospinal fluid for separation of the brain stem that was then rapidly frozen in optimal cutting temperature (OCT) medium for cryosectioning. The heart was removed and rinsed in cold PBS before being rapidly frozen in OCT. No more than ten minutes passed between decapitations and freezing the tissue in OCT for each animal.

The brain stems were sectioned at 10μm and collected on SuperFrost Plus glass slides in preparation for laser capture microdissection (LCM). Neurons in the DMV were identified by use of a rapid (∼15 minutes) immunofluorescence labeling procedure staining for NeuN. Pools of 5-10 neurons were captured using the Acturus HS Capsure system. Each cap containing neurons was visualized after capture to ensure that only neurons were included in the samples. Any cap containing non-neuronal cells were not used in the study. Apart from collecting neurons only, some samples were collected from the same slides that included the entire area of the DMV represented on that slice. The samples for this study were all collected from the part of the DMV that begins at the obex and runs caudally for 500μm.

### Gene expression assays

Pooled single cell samples were prepared for multiplex real-time quantitative polymerase chain reaction (RT-qPCR) with the Biomark HD system using VILO III (ThermoFisher) reverse transcription of RNA into cDNA followed by 22 cycles of pre-amplification (Taqman pre-amp master mix) of the cDNA using the collection of primers that were utilized in the multiplex RT-qPCR. Each set of primers was designed to span introns when possible and was validated through *in silico* PCR (Primer BLAST) and observation of a single band with the expected amplicon size on PCR gel using standard rat brain RNA, prepared for PCR by the same methods outlined previously. The quality of the results of the multiplex RT-qPCR was ensured through examination of the melt curves and standard curves. If there were multiple peaks for the melt curves or if a reliable standard curve was not present, the assay was not used in further downstream analysis. Similarly, if a sample showed that more than half of the assays were below the limit of detection, the entire sample was not utilized in downstream analysis.

The multiplex RT-qPCR experimental design included 84 genes, which are listed in Figure 2-1. The selection of these genes from the thousands expressed in the DMV was accomplished through inference based on our previous work in the dorsal vagal complex and through analysis of publicly available transcriptomic studies. To date, there is no publicly available transcriptomics data examining the DMV in the context of or any cardiovascular disease. There is, however, one dataset examining the DMV in human patients with Parkinson’s disease (PD) that may be helpful in understanding DMV pathology (GDS4154). Through use of ANOVA and template-matching, several genes that were differentially expressed in PD patients and only changed in the DMV were considered in constructing the gene list for this current work.

### Statistical Analysis

Raw C_t_ values from the multiplex RT-qPCR that passed melt curve based quality control were first median centered within each sample in order to account for the variations in total RNA in the sample to get delta C_t_ values. For analysis and reporting purposes, negative delta C_t_ (-dC_t_) values were used because a lower C_t_ value is representative of a higher expression level. By using -dC_t_ values, higher values now represent higher gene expression and are normalized to account for different initial total RNA amounts. In order to find statistical difference between treatments and over time, a mixed linear model ANOVA, aggregating the pooled single cell samples up to the animal level, was used with Tukey post-hoc analysis to correct for multiple comparisons. For determinations of gene expression correlation, each sample was taken to be independent and pairwise Pearson correlations between genes were computed. All statistical calculations were performed using R statistical software.

## Results

### Effects of cardiac ischemia on transcriptional state

Through the use of a hierarchical clustering algorithm with average linkage of Pearson correlations, it is possible to identify 6 sample clusters (SC) and 5 gene clusters (GC) through dendritic tree height cut-offs that parsimoniously segregate groups. This provides the framework for understanding the heterogeneous responses to cardiac ischemia over time (Figure 2). The distribution of samples from experimental groups among these 6 SC is not uniform as would be expected if LAD ligation did not alter the behavior of the DMV as a whole. Instead, a clear increase in the representation of certain transcriptional phenotypes with specific conditions and in many cases, distinct distributions of phenotypes on the left and right is observed (Figure 3). The 1 week sham (Sham-1) samples are generally divided into three main phenotypes, SC-A, SC-B, and SC-F. The 1 week LAD ligation samples (LAD-1) are distributed similarly, but with the additional representation in SC-C. The sham 3 week samples (Sham-3) are represented in all SC with negligible left/right differences. The 3 week LAD samples (LAD-3) distribute much differently than all of the other groups; SC-A and SC-F have only sparse representation whereas SC-C, SC-D, and SC-E are heavily represented, an effect that is exaggerated on the right side when compared to the left. All of these distributions are given in Figure 2 (pie charts above the heat map) and shown by treatment group in Figure 3A.

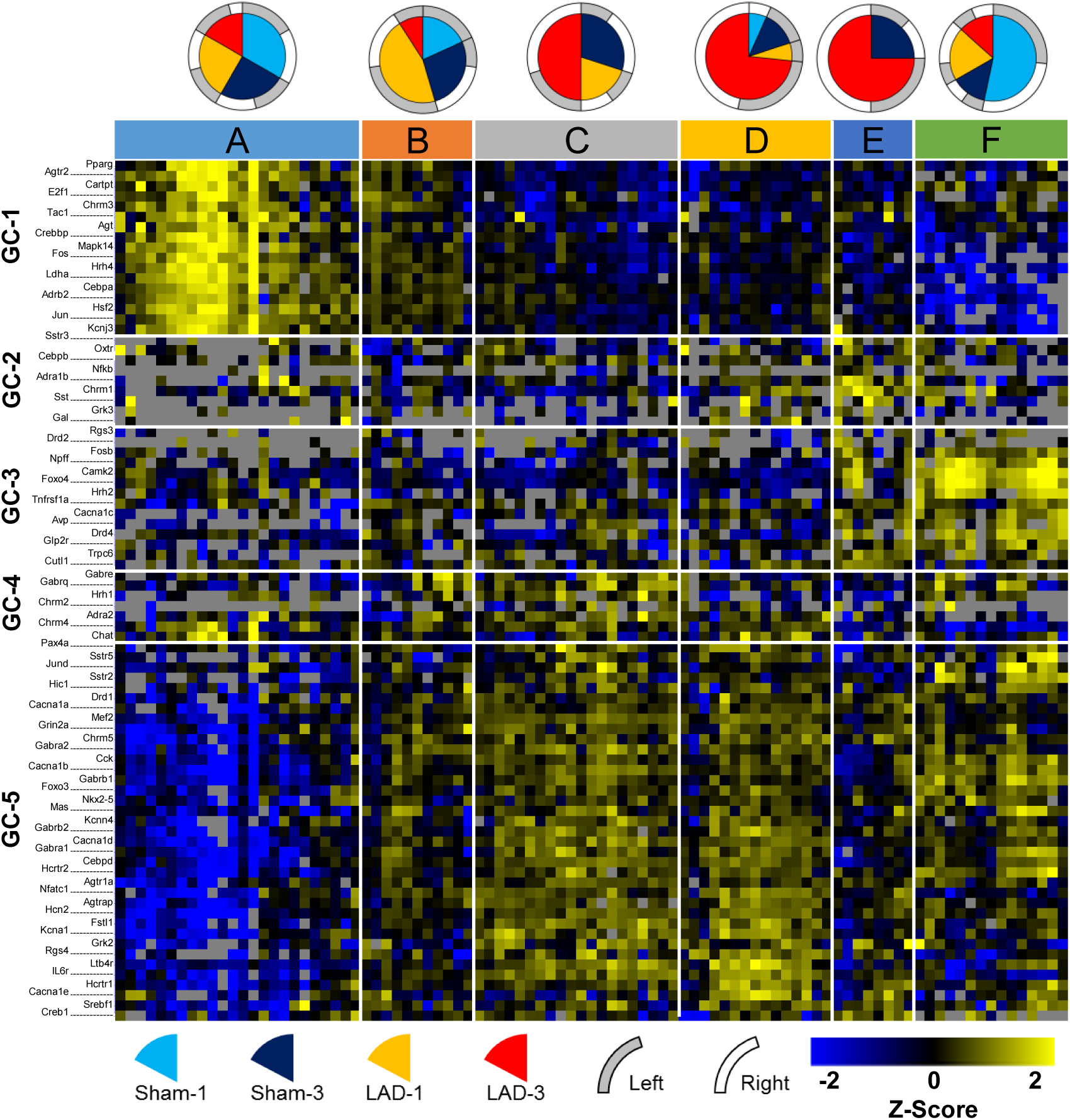

**Figure 3:**
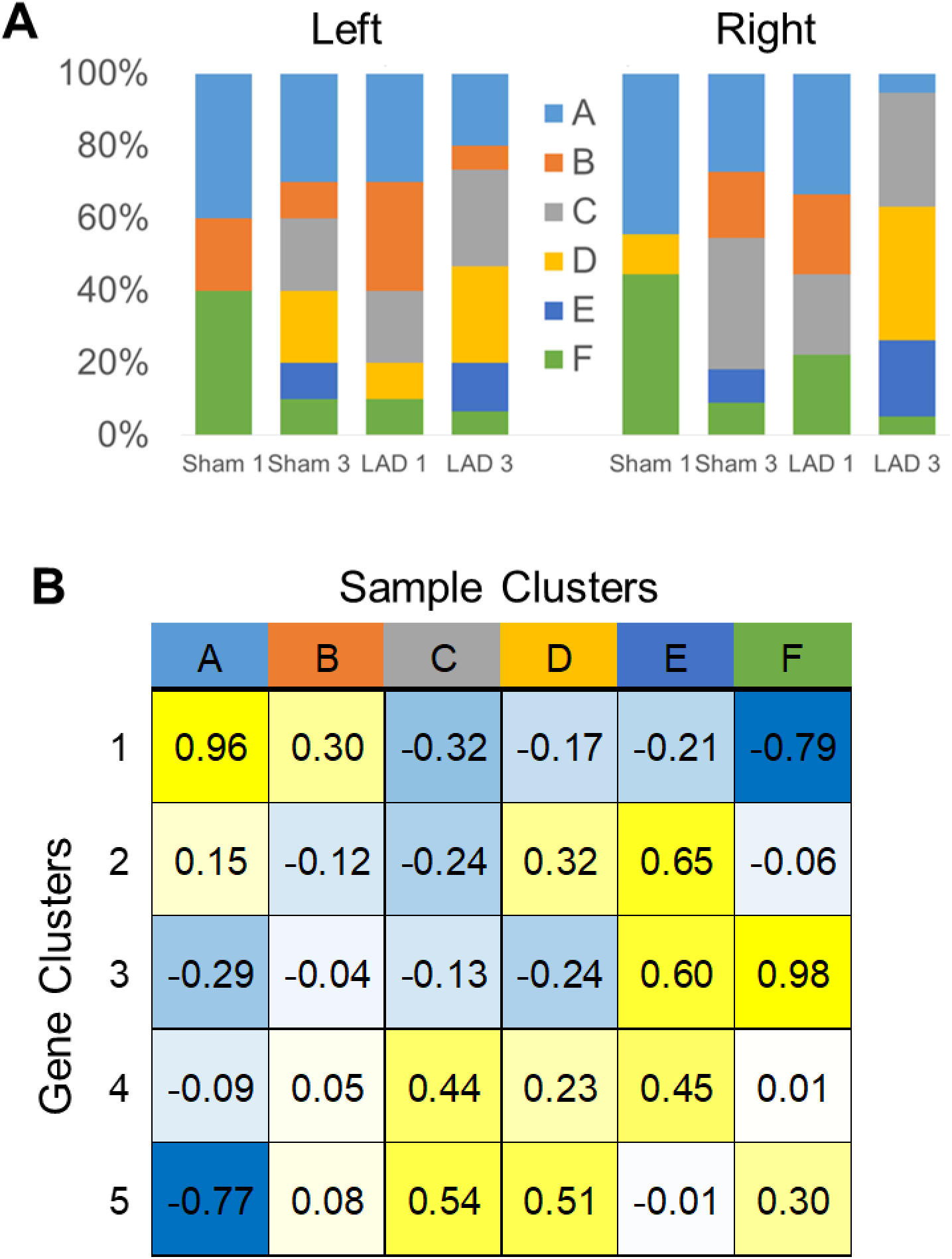
Summary information from heatmap hierarchical clustering in Figure 2. (A) Distributions for samples are segregated by their source from either the left or the right DMV. Sample cluster names and corresponding colors from the heatmap in Figure 1 are given in the legend in the center. (B) Average Z-score values for each sample cluster and gene cluster. Coloring is analogous to the heatmap in Figure 1 such that yellow is high expression and blue is low expression. Any *NA* values were excluded from averages.

These six sample clusters are generated generally from combinatorically differential expression across 5 gene clusters. The distinguishing features of SC-A are high expression of GC-1 and low expression of GC-5 with average levels of the others. SC-B may be considered as a transition from SC-A to SC-C with expression levels of nearly all of the GC in between those of SC-A and SC-C. SC-C has a characteristic high expression of GC-4 and GC-5 together and lower expression of GC-1 and GC-2. SC-D has high expression of GC-2 and GC-5 with more modest upregulation of GC-4 and downregulation of GC-3. SC-E includes upregulation of GC-2, GC-3, and GC-4 together. SC-F has a marked downregulation of GC-1 and marked upregulation of GC-3 with modest upregulation of GC-5. While not exact opposites, the composition of SC-A and SC-F have gene clusters (GC-1, GC-3, and GC-5) that have opposite expression tendencies. Of note as well is that both SC-A and SC-F are comprised of samples from all of the experimental groups, albeit in different proportions, perhaps suggesting their ubiquity in DMV function, irrespective of cardiovascular perturbation. Summaries of these characteristics can be seen in the aggregated Z-score heatmap of Figure 3B, derived from the heatmap in Figure 2.

The consequence of considering the pattern of gene expression through hierarchical clustering is that distinct functional groupings emerge that are differentially represented between treatment groups. The two sample clusters at the opposite ends of the heatmap in Figure 2, SC-A and SC-F, are the most clearly defined distinct transcriptional phenotypes based upon their average Euclidean distances. Clusters SC-B, SC-C, SC-D may initially all appear to be transitional groups that represent the populations of neurons that are in the process of shifting from SC-A to SC-F or vice versa. However, an examination of the minimum spanning tree (MST) in Figure 4A suggests that SC-B and SC-C are more likely transitional clusters whereas SC-D and SC-E are terminal phenotypes. The branching pattern for SC-E in the MST suggests that it is really two separate phenotypes that in this case differ due to ischemic cardiac damage. There is one branch that is comprised entirely of LAD-3 samples and has a distance of 3-4 in the MST from the bulk of the two main phenotypes, SC-A or SC-F. The other SC-E samples are from the Sham-3 group and are adjacent to the rest of the samples from SC-F.

**Figure 4:**
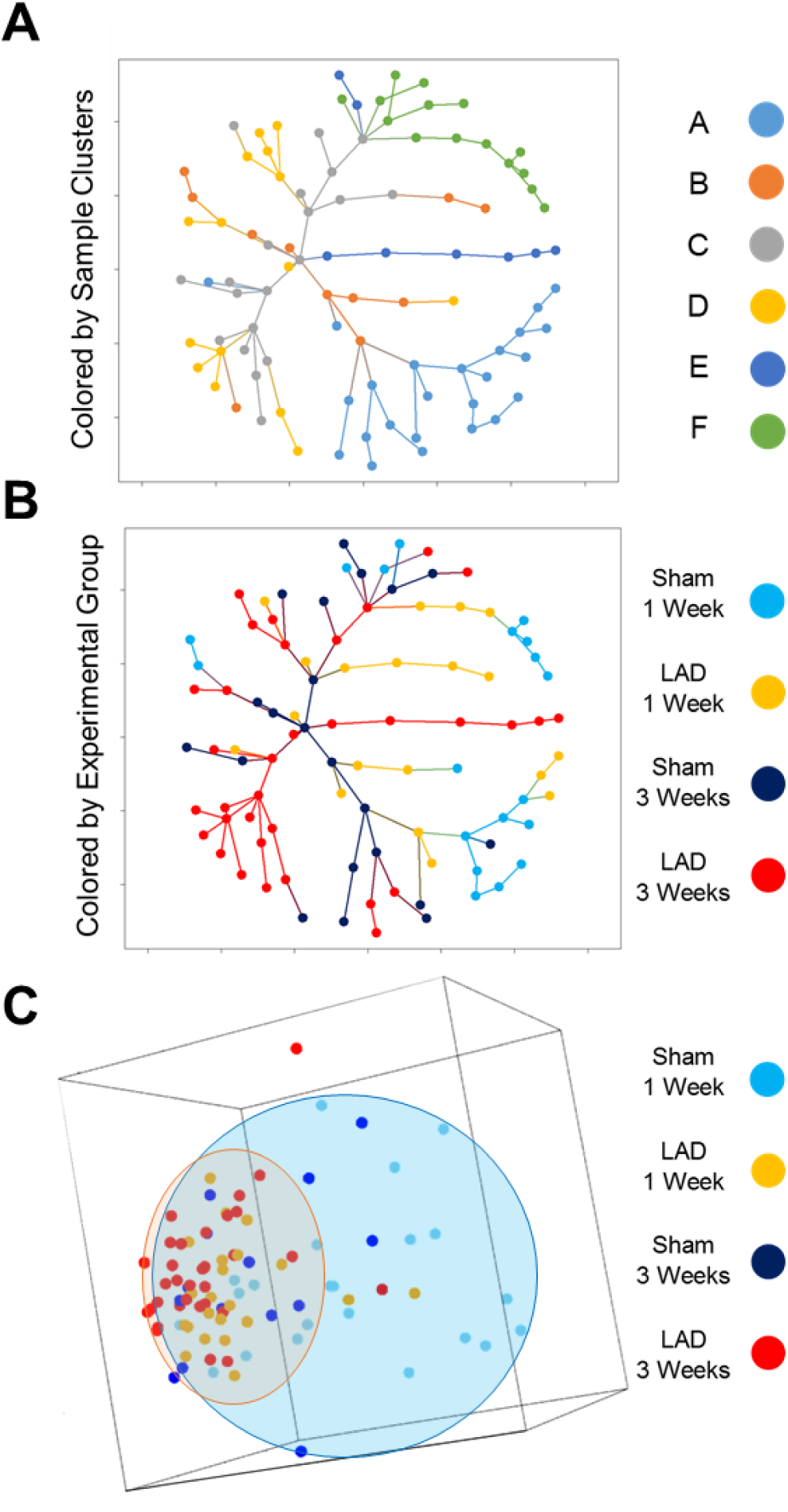
Dimensionality reduction though minimum spanning tree and principal component analysis demonstrates diminished expression space for LAD-3 samples toward the SC-C, SC-D, and SC-E expression patterns. (A) Minimum spanning tree colored by sample clusters shows that SC-B and SC-C are more transitional phenotypes with SC-A, SC-D, SC-E, and SC-F being more terminal. (B) The same minimum spanning tree as in (A), but colored for experimental groups, shows the unique terminal branches of LAD-3 comprised of samples of the SC-D and SC-E phenotypes. (C) Plot of first three principal components with ellisoids showing the general expression space for the sham samples (blue) and the LAD samples (orange) regardless of time point. Points are colored according to experimental group. This plot demonstrates a reduction in the expression space (diversity) of LAD samples toward an expression pattern that is present in some of the sham samples.

Consideration of the expression pattern across all measured genes using principal component analysis (Figure 4C) yields two interesting observations. The first is that the expression space, diversity of expression pattern, for the LAD ligation samples occupies a much more constrained space (red ellipsoid) than the sham samples (blue ellipsoid). The second observation of note is that this constrained space shown by nearly all LAD samples from both time points lies within the expression space defined by the sham samples. This suggests that there is a programmatic coordination in response to ischemic injury and that this response is within the realm of normal expression patterns at least for some neurons.. In this case the response phenotypes include SC-B and SC-C for LAD-1 and SC-C, SC-D, and SC-E for LAD-3.

### Effects of persistent cardiac ischemia on gene co-expression network topology

Gene co-expression networks were generated for each of the experimental groups using Pearson correlations and filtered using a q value cut-off of q<10^−3^. The unique edges for each of the networks generated are given in Figures 5 and 6 with the connectivity in any of the experimental groups were excluded from the full network for each group given in Extended Data section. Nodes (genes) with no network representations. It is clear from these network figures that the LAD-1 group has the greatest overall connectivity with over half of all edges being unique that group. Many of these unique edges are the negative correlation between genes in GC-1 and GC-5. Also of note are unique edges of interconnectivity within GC-1 and GC-5 respectively. The notable unique edges of the sham 1 week group include several negative correlations with Foxo4, Camk2, and Drd4 in GC-3 with genes in GC-1. Also, there are several unique positive correlations to genes in GC-5 from other GCs as well as some unique interconnectivity within GC-5. The LAD-3 group has much lower connectivity than either of the 1 week groups and also less unique edges. However, of interest is the emergence of Pax4a as a unique hub gene in its relationship to many GC-5 genes, the implications of which will be handled in the discussion that follows. The Sham-3 group is most notable for its very sparse connectivity.

**Figure 5:**
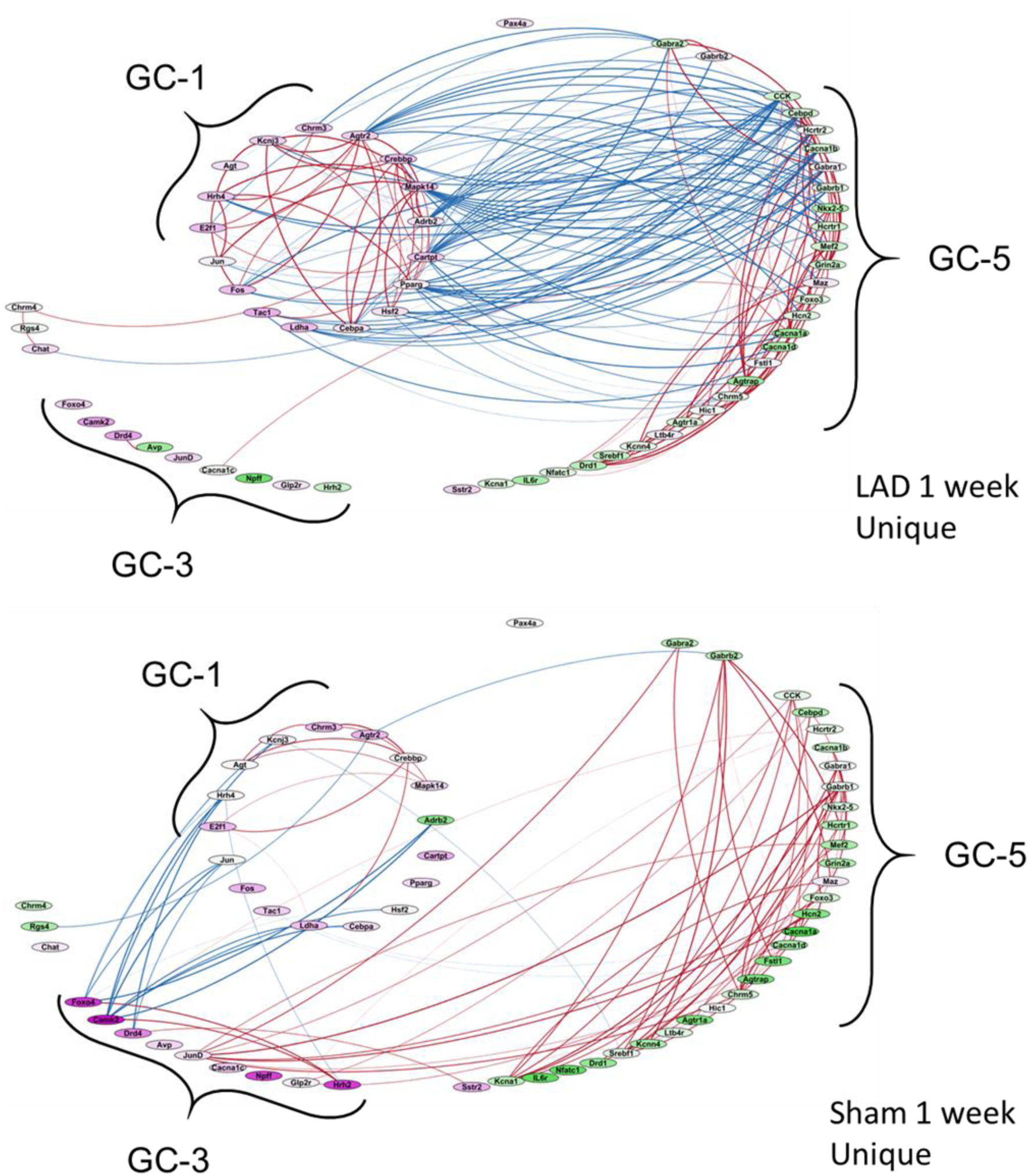
Unique gene co-expression networks for the 1 week time point of both experimental groups. Edges represent Pearson correlations with a q-value cut-off of q<10^−3^; only edges unique to a given experimental group are shown (full networks can be found in Extended Data).

**Figure 6:**
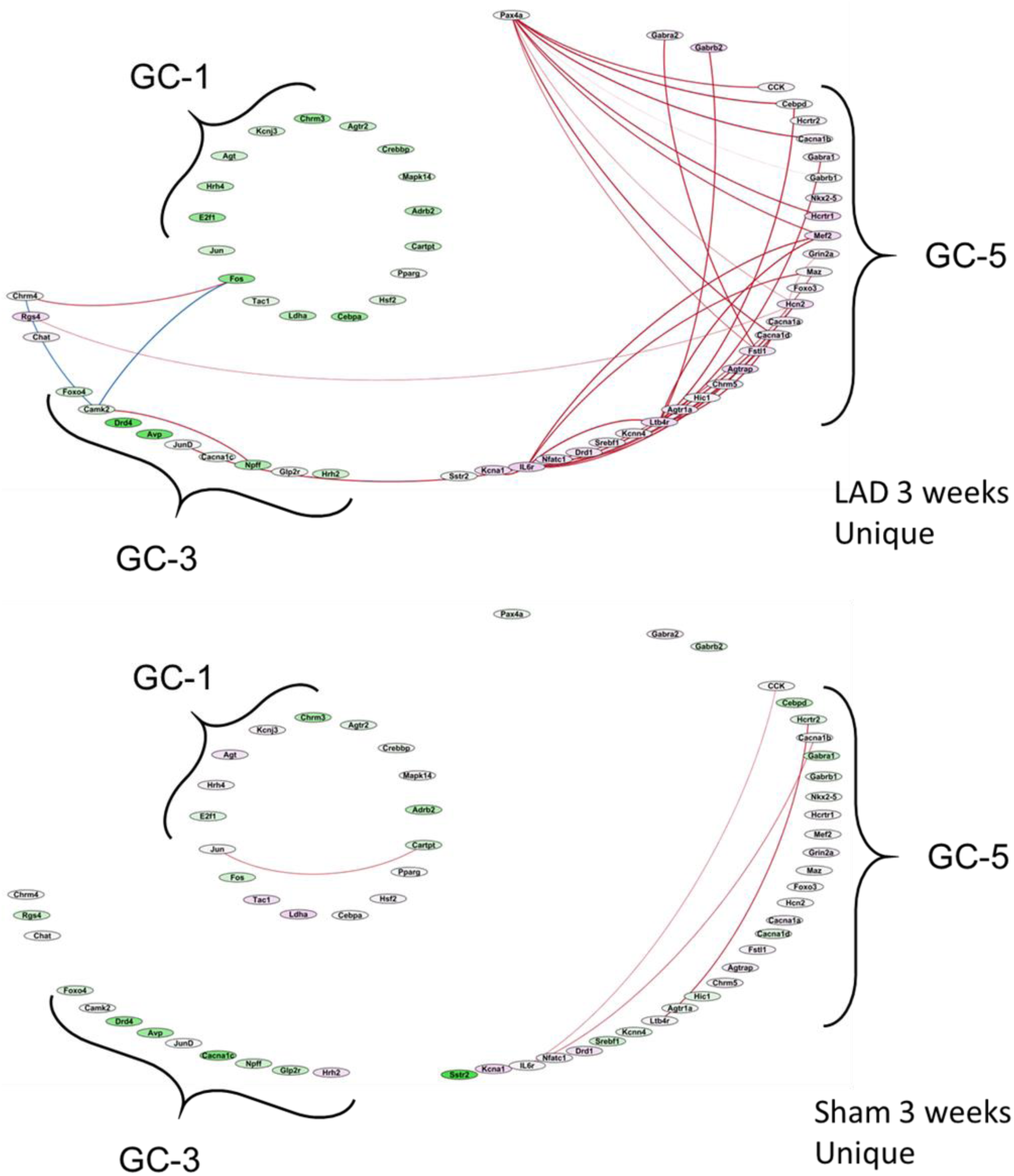
Unique gene co-expression networks for the 3 week time point of both experimental groups. Edges represent Pearson correlations with a q-value cut-off of q<10-3; only edges unique to a given experimental group are shown (full networks can be found in Extended Data).

### Effects of cardiac ischemia on individual gene expression

A differential expression analysis was performed using a nested ANOVA to account for multiple measurements within each animal. While several genes did show differential expression between the sham and LAD groups at both 1 week and 3 weeks post-surgery, these significant differences can be best explained by changes on either the right side or left side only rather than including all samples from both sides in aggregate. Of the six genes found to be significantly different, only Camk2 showed the bilateral difference of decreased expression in LAD-1 compared with Sham-1. This overall lower expression seems to be due to a loss of a bimodal distribution that is present in the sham condition. The other five genes from this study whose expression levels differed showed a pronounced effect on only the right or the left side (Figure 7). Only Sst (somatostatin precursor) was shown to change only on the left side of the DMV with the largely increased expression reaching significance at the 3 week time point. The right side did show increased Sst expression at 3 weeks as well, but it did not rise to the level of statistical significance due to the more broad distribution of expression values. Four genes showed significant changes on the right side of the DMV: Hrh2, Npff, Fosb, and Cacna1e. The largest effect size here is the diminished expression of Npff in LAD-1 compared with Sham-1. Fosb showed decreased expression on the right only at the 1 week time point and Cacna1e showed increased expression only at the 3 week time point. Hrh2 expression had an interesting pattern, wherein the expression on the right in the sham condition at both time points was much higher than the left, which was at a comparable level to both sides in the LAD condition at both time points. The LAD ligation surgery was associated with a shift down to left sided levels at both the 1 week and 3 week points. Considering these results through the lens of vagal projections to the gut that exhibit very distinct bilateral asymmetry (Fox and Powley 1985), it may be possible to interpret these changes in gene expression in a functionally relevant manner.

**Figure 7:**
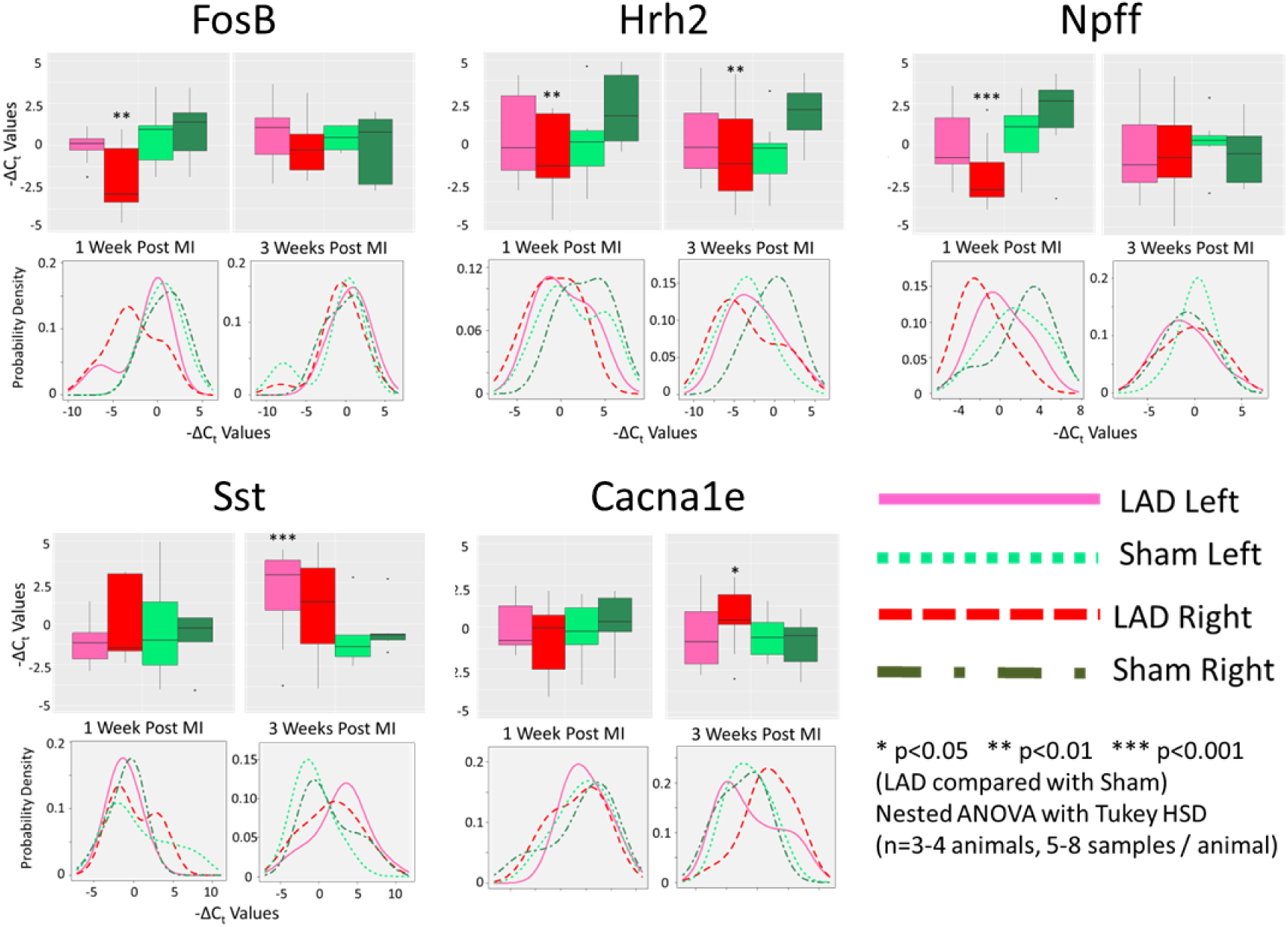
Genes with statistically different expression between experimental groups showed bilateral asymmetry. Of all the genes examined, only five showed statistically distinct expression levels between sham and LAD for either time point when accounting for animal level effects using a mixed linear model nested ANOVA (lme4 package in R). For each gene, boxplots of gene expression values are shown broken down by quartiles. Asterisks (*) indicate statistically significant differences between the sham and LAD surgery groups within a given time point on either the left or right side (as indicated by box/line color). Also shown for each gene is a density plot for the expression values for both left/right and sham/LAD conditions within a time point.

**Figure 8:**
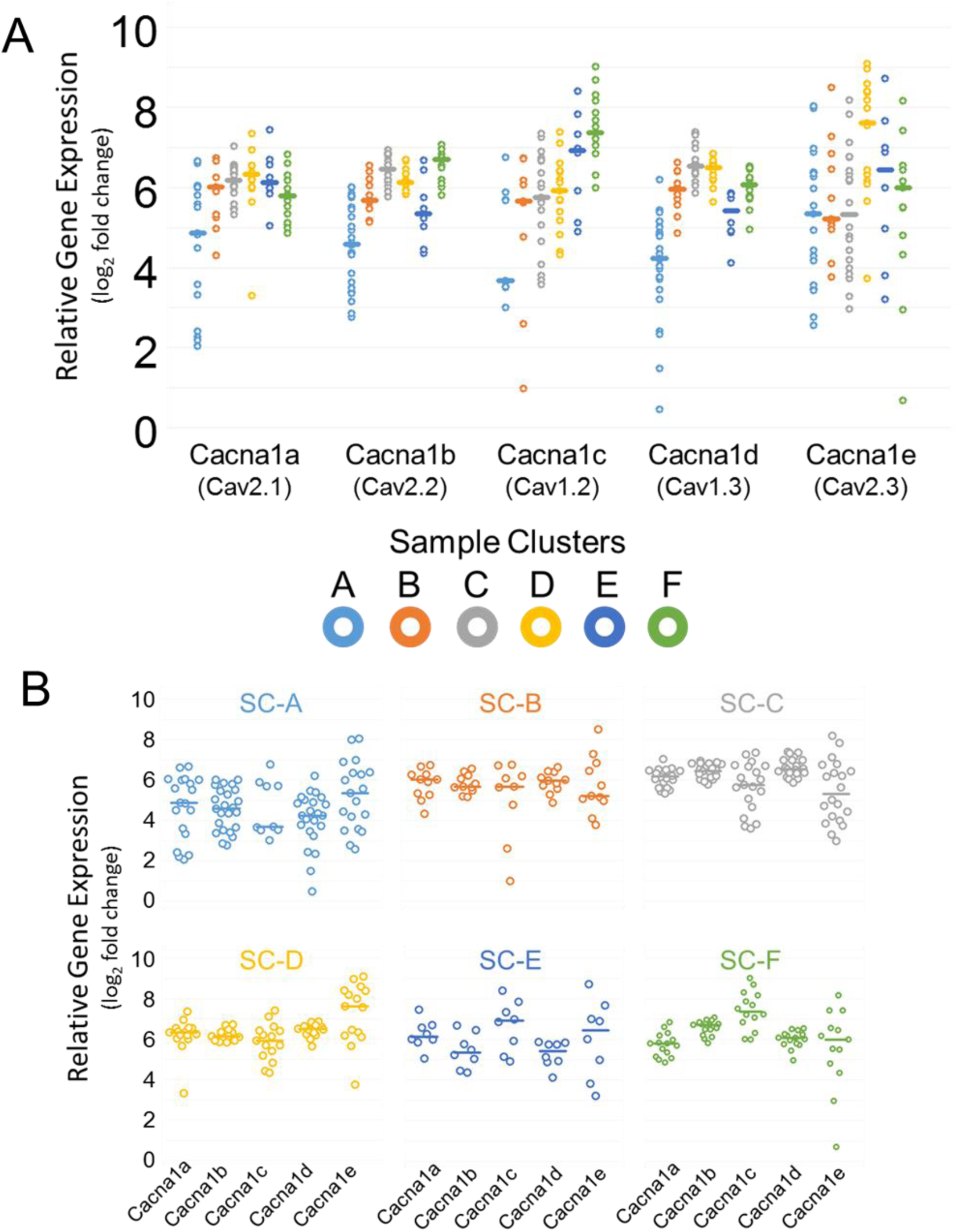
Expression of voltage gated calcium channels across samples clusters show some distinct patterns that may hint at functional behavior. (A) Each calcium channel’s gene expression values across all sample clusters with each dot representing one sample of pooled neurons. (B) Each sample cluster’s pattern of calcium channel expression with each dot representing a pooled neuron sample. Both (A) and (B) show the same data points, only grouped differently to aid in making comparisons.

## Discussion

### Ischemic heart failure response phenotype

There appears to be at least one ischemic heart failure response phenotypes for neurons in the DMV, as suggested by the uniquely large over-representation of LAD-3 samples in SC-C, SC-D, and SC-E. Of note is the upregulation and unique connectivity of the transcription factor Pax4a, a transcription factor is strongly regulated by the transcriptional repressor REST which drives stem cells into a neurosecretory phenotype (Bruce et al. 2006; Kemp et al. 2003). The disinhibition of gene expression through the downregulation of REST with concomitant upregulation of Pax4a is restricted largely to neurons, but is also a feature of developing/regenerating pancreatic islet cells, in particular β and δ cells (Biason-Lauber *et al.*, 2005). The coordination of increased expression of the genes in GC-5 may be accomplished in part through effects of Pax4a as evidenced by its unique network connectivity in that condition (Figure 5). Coupled with the coordinated upregulation of Cacna1b, Cacna1d, Cck, Gabra1, and Gabra2 the pictures is painted less of a neuron and more of an enteroendocrine cell, like a pancreatic β/δ cells or colonic L/I cells (Gameiro et al. 2005; Rogers et al. 2011; Schonhoff et al. 2004). This is interesting given that the canonical satiety peptides (CCK, GLP-1, SST, etc) can play a protective effect in the face of ischemic damage at the heart (Sartor and Verberne 2010; Mukharji et al. 2013; Grossini et al. 2013; Goetze et al. 2016; Dong et al. 2017; Mobley et al. 2006).

A more broad interpretation of the main neuronal phenotypes considering all genes examined here suggests that SC-A and SC-F are the phenotypes of DMV neurons that exist either at resting state or are recruited for generalized injury/stress response. This is shown in the distribution of sample clusters within experimental groups in Figure 3A. The combination of nearly equal representation of all experimental groups for each and the distinct gene expression patterns drive this conjecture. While the heatmap makes SC-C and SC-D appear very close in expression patterns, examination of the spanning tree suggests that SC-D is more of a terminal phenotype whereas SC-C is the plastic, transitional phenotype through which neurons must transition before moving from one of the other phenotypes. Given the poorly defined separation of these clusters, the most likely case is that they are the same phenotype, but are behaving in different ways, perhaps a result of inputs or feedback, but are operating under the same general genetic program. The main difference between SC-C and SC-D is the higher expression of GC-2 in SC-D, a gene module that contains neuropeptides like somatostatin and galanin along with the somatostatin-3 receptor and the oxytocin receptor. It also has a clear upregulation of the beta-1 adrenergic receptor, better known for its cardiac specificity in the periphery, and for which little is known about function in neurons of vagal motor neurons. Secondarily is the upregulation of GC-4 in SC-C compared with SC-D. GC-4 contains several inhibitory receptors: two GABA receptor subunits (ε and θ), two G_i_ coupled muscarinic receptors, and the G_i_ coupled alpha 2 adrenergic receptor. Coupled with the relative increased expression of several calcium channel genes (save for Cacna1c), this is suggestive of a neuronal phenotype capable of pacemaker or even burst firing (Cui et al. 2018; Yang et al. 2018; Pape and McCormick 1989; Zhu et al. 1999). Also contributory to this conjecture is the upregulation of Grin2a, Hcn2, and Kcnn4, all of which are involved in autonomous firing (Dong et al. 2016; Chan et al. 2004; Yang et al. 2018). Along with the unique inhibitory receptors from GC-4 there is also increased expression of Gabra1 and Gabrb1 GABA receptor subunits, which may be necessary to lower the resting membrane potential of these neurons or prevent large depolarization events to permit their function as autonomous pacemakers capable of burst firing (Cui et al. 2018; Yang et al. 2018; Zhu et al. 1999). Also of interest in SC-C and SC-D is the increased expression of several receptors that are responsive to appetite-regulating hormones including two somatostatin receptor subtypes and both orexin receptor subtypes. This may be indicative of the type of signaling these projections are involved with at the gut.

### Differences in network topology

Within each of the treatment cohorts, there are several highly connected genes as is the baseline expectation for a biological network that is suspected to have scale-free topology (Barabási 2009). Overall, there few genes that are highly connected across most of the cohorts, most notable being Cacna1b that is a member of the top five connected genes in each cohort. Consideration of the unique hubs that emerge is perhaps more telling (obtained by only considering the edges that are unique to a given cohort). Cebpd is highly connected in the LAD-1 group and the LAD-3 group, but with many unique edges in the LAD-1 group. This suggests a role of Cepbd in mediating gene expression in the acute response to cardiovascular injury that persists to some extent beyond the acute phase. There is a relatively high connectivity for Cebpd across some of the Sham cohorts as well, but not nearly to the extent that exists in the LAD cohorts. Interestingly, Cebpd expression levels in the LAD-1 group are significantly different, but that there is such disparate connectivity may point to differential activation states of the protein product of the gene, differing efficiency of translation, or something of the like. While the work on Cepbd in neurons is limited, in neuron-like pancreatic beta cells Cebpd mediates an anti-apoptotic and anti-inflammatory state (Moore et al. 2012).

Although the connectivity among the measured genes in the LAD-3 group is not as high as the either of the 1 week groups, there are still some inferences to be made from what is uniquely connected. There are three genes of interest here: IL6r, Ltb4r, and Pax4a. While IL-6 is a well-described pro-inflammatory cytokine, this is not to suppose that the effects of IL-6 binding its receptor in all contexts serves to increase inflammation. In neurons, the IL-6 receptor mediates anti-apoptotic signaling and can serve to protect against reactive oxygen species associated with metabolic stressors (Thier et al. 1999; März et al. 1998). The levels of IL6r do not differ significantly between LAD-1, LAD-3 and Sham-3 (it is suppressed in Sham-1 relative to the others). However, high connectivity of IL6r in the LAD-3 group suggests that there is some signaling that mediated by the protein product that is having an effect on expression of other genes, including Ltb4r and the pacemaker contributing channels Kcna1 and Hcn2. Similarly, the leukotriene B4 receptor is a canonical mediator of inflammatory processes, but has a different role in neurons of the central nervous system, promoting neurogenesis and neural differentiation from progenitor stem cells (Wada et al. 2006). Interestingly, there is a great deal of evidence for a similar role for the IL-6 receptor in neurons of the CNS (Islam et al. 2009). It may be possible that cohort of genes associated with IL6r and Ltb4r expression are related to neural progenitor phenotype if stem cells were under consideration. However, these cells are neurons and were selected based upon strong NeuN protein expression as evidenced through immunofluorescence. Since NeuN is only expressed in neural progenitor cells after the neuron “fate” has been determined (Gusel’nikova and Korzhevskiy 2015), it is highly unlikely that any stem cells of pre-neuronal phenotype were selected for this study if any do in fact exist in the DMV of adult rats. Therefore, it is possible that reactivation of some of the differentiating cell programming is used to regress the terminally differentiated neurons and permit them to undergo a phenotype shift. The third unique hub gene may give a clue to the nature of this phenotype shift, since Pax4a mediates differentiation into a neurosecretory phenotype both in the central nervous system as well as for the specialized secretory cells in the pancreas (Schonhoff et al. 2004; Biason-Lauber et al. 2005). This is supported by upregulation of several genes in GC-5 that are coordinated by the three hub genes, IL6r, Ltb4r, and Pax4a. Many genes in GC-5 are suggestive of a neurosecretory phenotype, including several ion channels (Cacna1d, Kcna1, Hcn2) (Goldberg et al. 2012; Cooper et al. 2015), neuropeptide signaling (Sstr2, Sstr5, Cck, Hcrtr1, Hcrtr2), and GABA receptors (Gabra1, Gabra2, Gabrb1, Gabrb2) (Gameiro et al. 2005). While not verified directly in this work here, many of the genes that are well-connected n the unique LAD-3 network are those under the influence of the repressor element 1 silencing transcription factor (REST) complex, an essential regulator of a neurosecretory phenotype (Bruce et al. 2006; Mieda et al. 1997; Wood et al. 1996). More work is needed to consider the possibility that repression of REST plays a role in the phenotype shift toward a neurosecretory phenotype.

## Conclusion

In response to ischemic heart failure, neurons of the dorsal motor nucleus undergo a transcriptional shift towards a neurosecretory phenotype. This was evidenced by the upregulation of ion channels associated with a pacemaker phenotype, an upregulation several neuropeptides, and increased expression and influence of Pax4a. This response to chronic ischemic damage in the heart demonstrates that neurons of the DMV are capable of adaptive responses in the context of peripheral organ damage. More work will be needed to understand if these changes are beneficial or harmful to the cardiovascular system, particularly with regard to the time course. Changes are beneficial early on in the adaptive process to injury may prove to have deleterious trade-offs in the end stages of the injury process. Regardless, the DMV and its vagal efferent projections are adaptive to peripheral injury and those implications warrant further investigation as to how this effects the rest of the body system and whether such adaptions can give clues to novel therapeutic intervention for conditions like cardiac ischemia.

---

**Figure 2-1:**
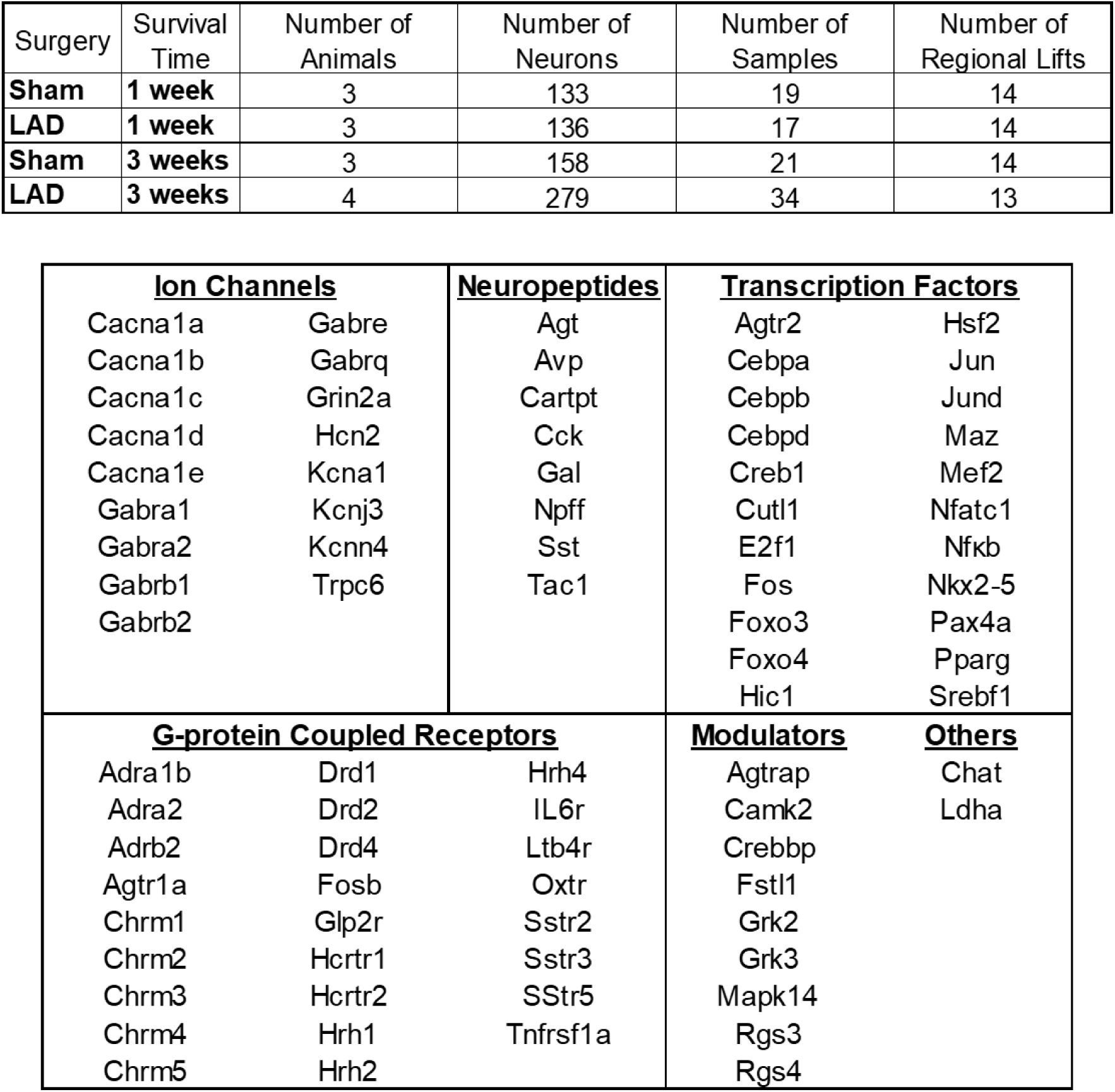
Tables of samples and gene assays. Samples were derived from cell pools of between 5-10 cells. See Appendix A for gene selection rationale and Appendix B for primer sequences used in multiplex RT-qPCR.

**Figure 5-1:**
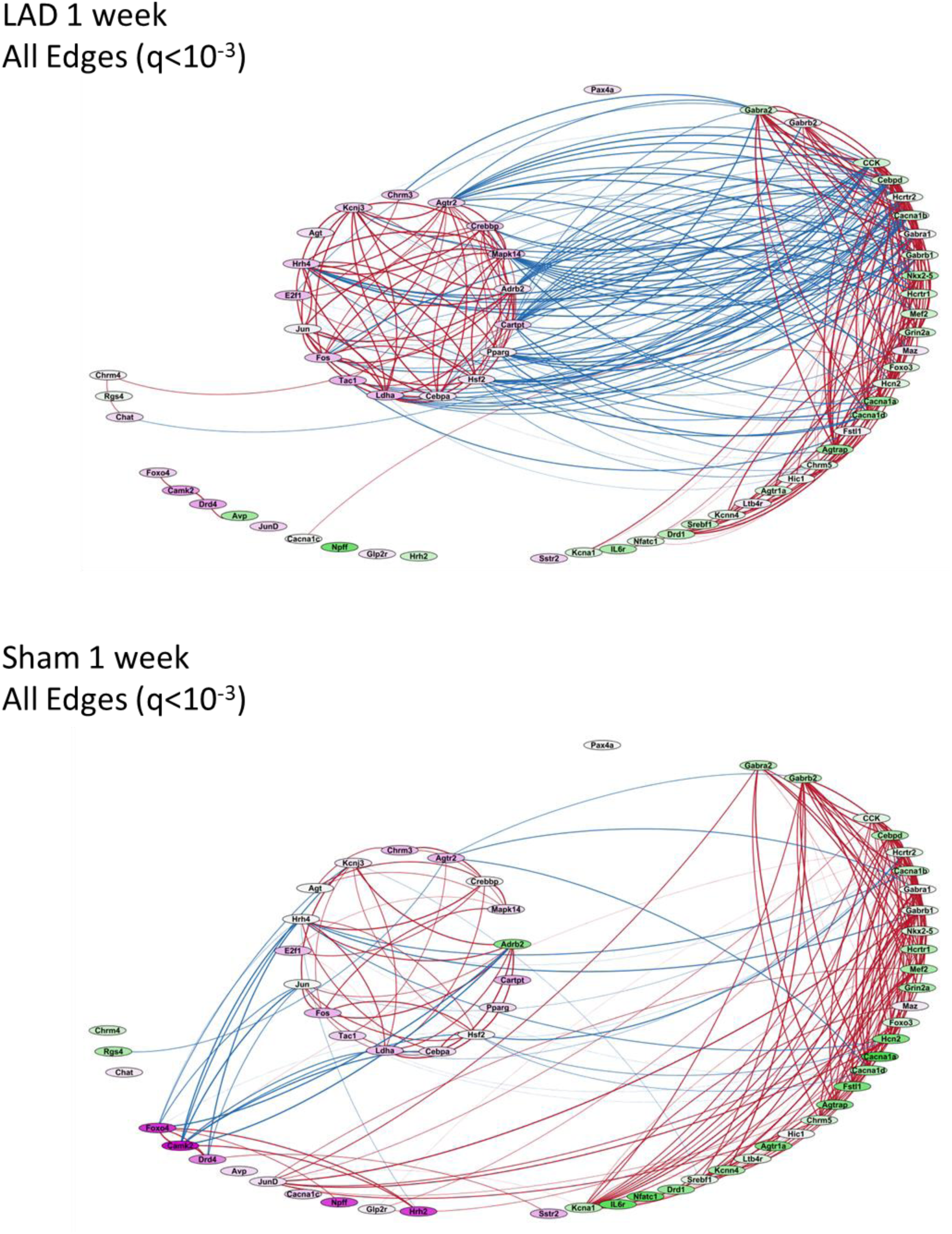
Full networks for LAD and Sham 1 week cohorts. Nodes are genes and edges are correlations between two genes. Green nodes have low expression and violet nodes have high expression with a 32-fold difference in between their relative expression. Blue edges are negative correlations and red edges positive with the edge thickness proportional to the correlation coefficient.

**Figure 6-1:**
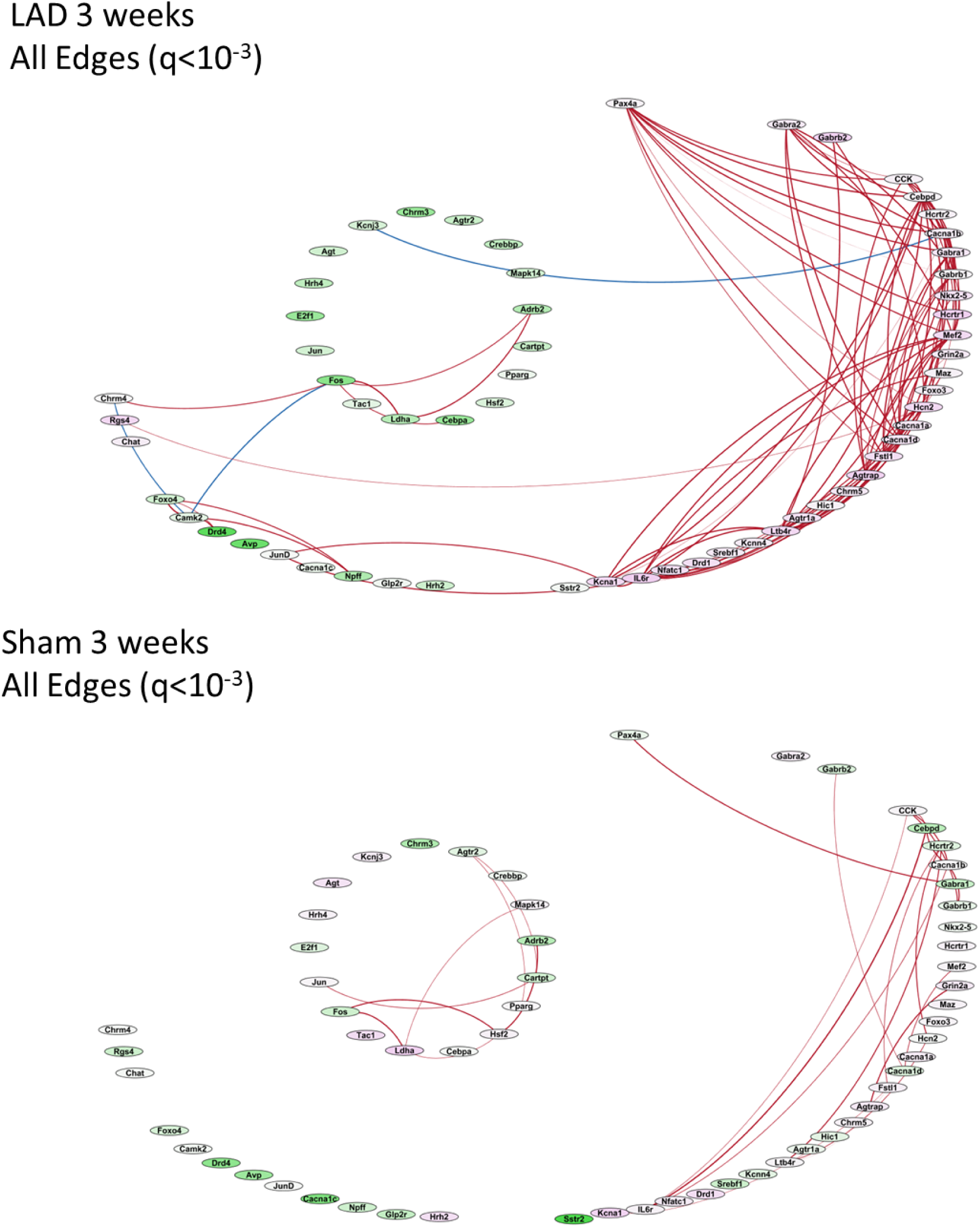
Full networks for LAD and Sham 3 week cohorts. Nodes are genes and edges are correlations between two genes. Green nodes have low expression and violet nodes have high expression with a 32-fold difference in between their relative expression. Blue edges are negative correlations and red edges positive with the edge thickness proportional to the correlation coefficient.

**Figure 6-2:**
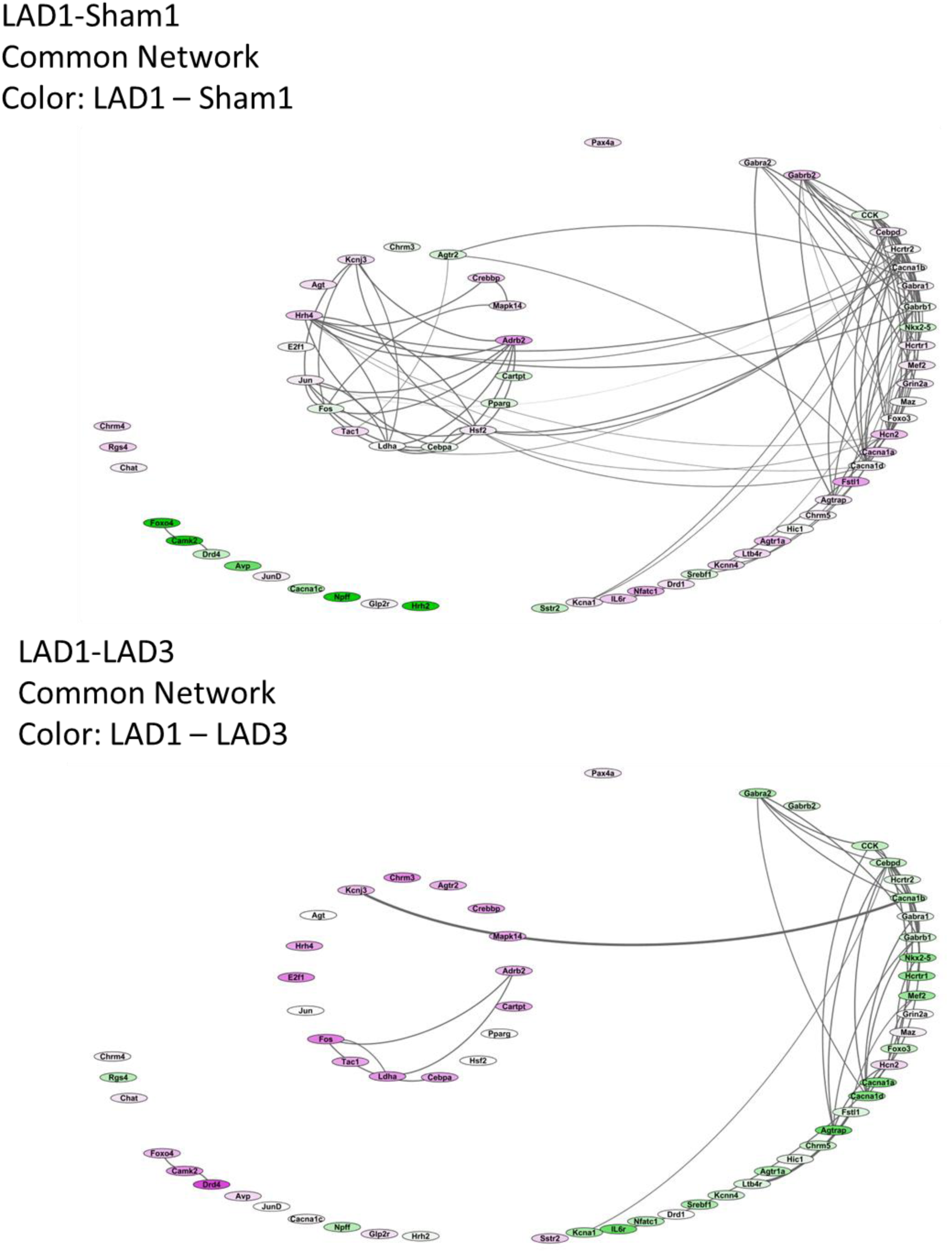
Network representations of common edges between two groups. Edge thickness is proportional to the average correlation coefficient. Green nodes indicate gene for which the expression of the LAD-1 group was lower than the comparison and violet nodes indicate genes for which the LAD-1 group was higher in expression. There is a 32-fold difference in gene expression between extreme green and extreme violet colors.

## Acknowledgements

The authors would like to thank Dr. Julian Paton and Dr. Angelo Lepore for their questions and input during the development of this work.

This work was made possible by National Institutes of Health grants: U01 HL133360; R01 HL111621-01A1; OT2 OD023848

## Notes

None of the authors have any conflicts of interest to disclose.

